# Sympatry with the devil? Niche-based approach explains the mechanisms allowing coexistence of native and non-native salmonids

**DOI:** 10.1101/068171

**Authors:** Camille Musseau, Simone Vincenzi, Dušan Jesenšek, Stéphanie Boulêtreau, Frédéric Santoul, Alain J Crivelli

**Affiliations:** Department of Biology, Chemistry, Pharmacy, Institute of Biology, Freie Universität Berlin, Königin-Luise-Strasse 1 – 3, Berlin 14195, Germany; Leibniz-Institute of Freshwater Ecology and Inland Fisheries (IGB), Müggelseedamm 310, 12587 Berlin, Germany; Ecolab, Université de Toulouse, CNRS, INPT, UPS, 118 route de Narbonne, 31062 Toulouse, France; Center for Stock Assessment Research, Department of Applied Mathematics and Statistics, University of California, Santa Cruz, Santa Cruz, California, United States of America; Dipartimento di Elettronica, Informazione e Bioingegneria Politecnico di Milano, Milan, Italy; Tolmin Angling Association, Modrej 26a, 65216 Most na Soci, Slovenia; Station Biologique de la Tour du Valat, Le Sambuc, 13200 Arles, France

**Keywords:** stable isotopes, niche shift, niche replacement, niche overlap, rainbow trout, marble trout

## Abstract

Niche-based hypotheses have been proposed to explain processes and mechanisms of success in the establishment of non-native species into native communities. Competition due to niche overlap may lead to native species niche shift and to native species replacement. To understand the ecological consequences of trophic interactions between non-native rainbow trout and native and endangered marble trout, we used as model system the Idrijca river (Western Slovenia) in which marble trout occurs either in allopatry (MTa) or in sympatry (MTs) with rainbow trout (RTs). We focused on different metrics of niche change such as centroid shift, niche overlap and trophic niche breadth using stable isotope analysis (δ^15^N and δ^13^C). Our results showed plasticity in niche overlap between MTs and RTs and niche shift of marble trout when occurring in sympatry with RTs, but not due to a niche replacement of MTs by RTs. Niche breadth of marble trout increases in sympatry and the trophic position during the growth period was higher for MTs than MTa.

## Introduction

Biological invasions are among the main causes of the current biodiversity decline (Mack et al., 2000; Clavero & García-Berthou, 2005). Different niche-based hypotheses have been proposed to explain mechanisms underlying successful biological invasions leading to coexistence of species occupying a similar ecological niche or species replacement, along with the ecological impacts of biological invasions (Ricciardi et al., 2013). The use of unexploited resources (i.e. ‘empty niche’ or ‘unfilled niche’) by non-native species may reduce competition with native communities and enable the coexistence of species occupying a similar niche in the same habitat (Schoener, 1974; Chesson, 2000; Okabe & Agetsuma, 2007; Mason et al., 2008; Juncos et al., 2015). On the other hand, non-native species that are superior competitor can compete for the same resources used by the native species. Competitive exclusion can lead to niche shift for the native species and/or a niche replacement of the native species, thus leading through reduction of food intake for the native species to slower individual growth, lower population density, and possibly species extinction (Bøhn et al., 2008). The fundamental niche of a species defines its ecological properties without competitive interaction with other species, and is theoretically expected to be broader than the realised niche in the presence of interspecific interactions (Hutchinson, 1957; Pearman et al., 2008; Holt, 2009). In the context of biological invasions, some studies have shown a realised niche smaller than fundamental niche for native species due to competitive interactions between native and non-native species for space or food (Wauters & Gurnell, 1999; Hamidan et al., 2015).

The trophic niche characterizes the functional role of organisms in a food web, as it reflects both the diversity of resources used by a consumer and the trophic interactions in the system. Since the trophic niche of native species can be affected by non-native species through competition for the same food resources, it is fundamental to investigate and understand the effects of non-native species on food webs of native recipient ecosystems (Sagouis et al., 2015). In recent years, the use of stable isotope niches as a proxy of trophic niches has gained popularity (Newsome et al., 2007, 2012; Layman et al., 2012) also thanks to the development of community-wide measures and statistical methods to characterize trophic niches (Layman et al., 2007; Turner et al., 2010; Jackson et al., 2011).

Many non-native fish species have established self-sustaining populations in European freshwaters (Strayer, 2010). Especially in headwater streams, non-native salmonids such as rainbow trout in Europe and brown trout in the US have been stocked for angling (Cucherousset & Olden, 2011). Invasive salmonids are often detrimental to native salmonid populations because of potential predation and competition for food and space (Hasegawa et al., 2004; Morita et al., 2004) and are a serious threat to the persistence of native populations (Krueger & May, 1991; Takami et al., 2002; Seiler & Keeley, 2009; Hasegawa et al., 2014). Invasion by non-native salmonid often lead to reduced growth (Carlson et al., 2007; Van Zwol et al., 2012; Houde et al., 2015a), survival (Blanchet et al., 2007; Houde et al., 2015b) and lower abundance of the native species than that of the invasive species (Nomoto et al., 2010; Benjamin & Baxter, 2012). In some cases, the native salmonid species persist at higher abundances than the non-native species, which suggests a segregation of ecological niches, such as different habitat occupied or food consumed (Inoue et al., 2009; Korsu et al., 2009; Hasegawa et al., 2010, 2012).

Rainbow trout (*Oncorhynchus mykiss*) has been introduced worldwide and has been listed among the worst invasive species by the IUCN (International Union for Conservation of Nature, Lowe et al., 2000). Numerous studies showed substantial negative effects (i.e. hybridisation, disease transmission, predation and competition) of non-native rainbow trout on native salmonids in a large geographical range, including Europe (Blanchet et al., 2007; Stanković et al., 2015). Rainbow trout have been introduced in Slovenia in early 20^th^ century for recreational purposes and have established several self-sustaining populations (Stanković et al., 2015). One of these introductions occurred in 1962 in the upper part of the Idrijca River. The rainbow trout population of the Idrijca River (RTs) has been self-sustaining following the introduction and lives in sympatry with the endemic and endangered marble trout (*Salmo marmoratus*). A population of marble trout lives in allopatry in the upper reaches of the same river. Previous work showed minor effects of rainbow trout on body growth and survival of marble trout in sympatry (MTs) when comparing those traits to those of marble trout in allopatry (MTa), and showed stable species coexistence between the non-native and native species (Vincenzi et al., 2011), with marble trout in sympatry older than juveniles that are on average 3 times as abundant as rainbow trout.

Evolutionary models of community assembly often assume that coexistence results from the operation of resource partitioning mechanisms that competition among similar species (Macarthur & Levins, 1967). Our goal is to investigate the trophic niche based-mechanisms that allow long-term coexistence of the two salmonid species using stable isotope methods. Specifically, we hypothesise (i) niche segregation between MTs and RTs; (ii) no displacement of the marble trout’s niche; (iii) no niche breadth difference between MTa and MTs and (iv) no niche replacement of the native species by rainbow trout.

## Material and Methods

### Study area

Sampling was conducted in 2012 (June) and 2013 (June and September) in the Idrijca River (South-Western Slovenia). The Idrijca River is a 60km-long tributary of the Soča River. The upper part of Idrijca watershed is mainly covered by deciduous forests (*Fagus sylvatica*), with low human activity. A dam in Idrijca prevents fish movement from downstream to upstream and provides two sectors of interest, Upper and Lower Idrijca. In Upper Idrijca marble trout live in allopatry (Berrebi et al., 2000; Fumagalli et al., 2002). In Lower Idrijca, marble trout live in sympatry with rainbow trout (Vincenzi et al., 2011). Marble trout (Upper and Lower Idrijca) and rainbow trout (Lower Idrijca) are the only fish species living in those sectors.

The two sectors are separated by approximately 1 km. The altitude ranges are 718-720 m above sea level (first sector) and 537-543 m (second sector). Pools occupy 65% of sector total surface of Upper Idrijca and 40% of the sector total surface of Lower Idrijca. Annual mean temperatures were similar in the two sectors in 2012 (allopatry: 7.42°C ± 3.14 and sympatry: 7.94°C ± 3.51) and in 2013 (allopatry: 7.87°C ± 3.11 and sympatry: 8.39°C ± 3.42).

### Fish and invertebrates sampling

Sampling surveys were carried out on the whole length of each station starting downstream using a gasoline-powered portable backpack electrofishing unit. Each station session was electrofished two times, allowing to produce a multiple-pass removal estimate of trout abundance using Microfish 3.0 (Van Deventer & Platts, 1989). More information on the study site can be found in Vincenzi et al. (2011) and densities in ind·ha^-1^ estimated in June and September from 2002 to 2014 for MTa, MTs and RTs are reported in Fig.1. We collected fin tissue for the stable isotope analysis (as a proxy of muscle isotopic composition, Jardine et al., 2005) in 2012 and 2013 from 633 fish: 246 from marble trout in allopatry (MTa), 224 from marble trout in sympatry (MTs) and 163 from rainbow trout in sympatry (RTs) (Table 1).

**Fig. 1.**
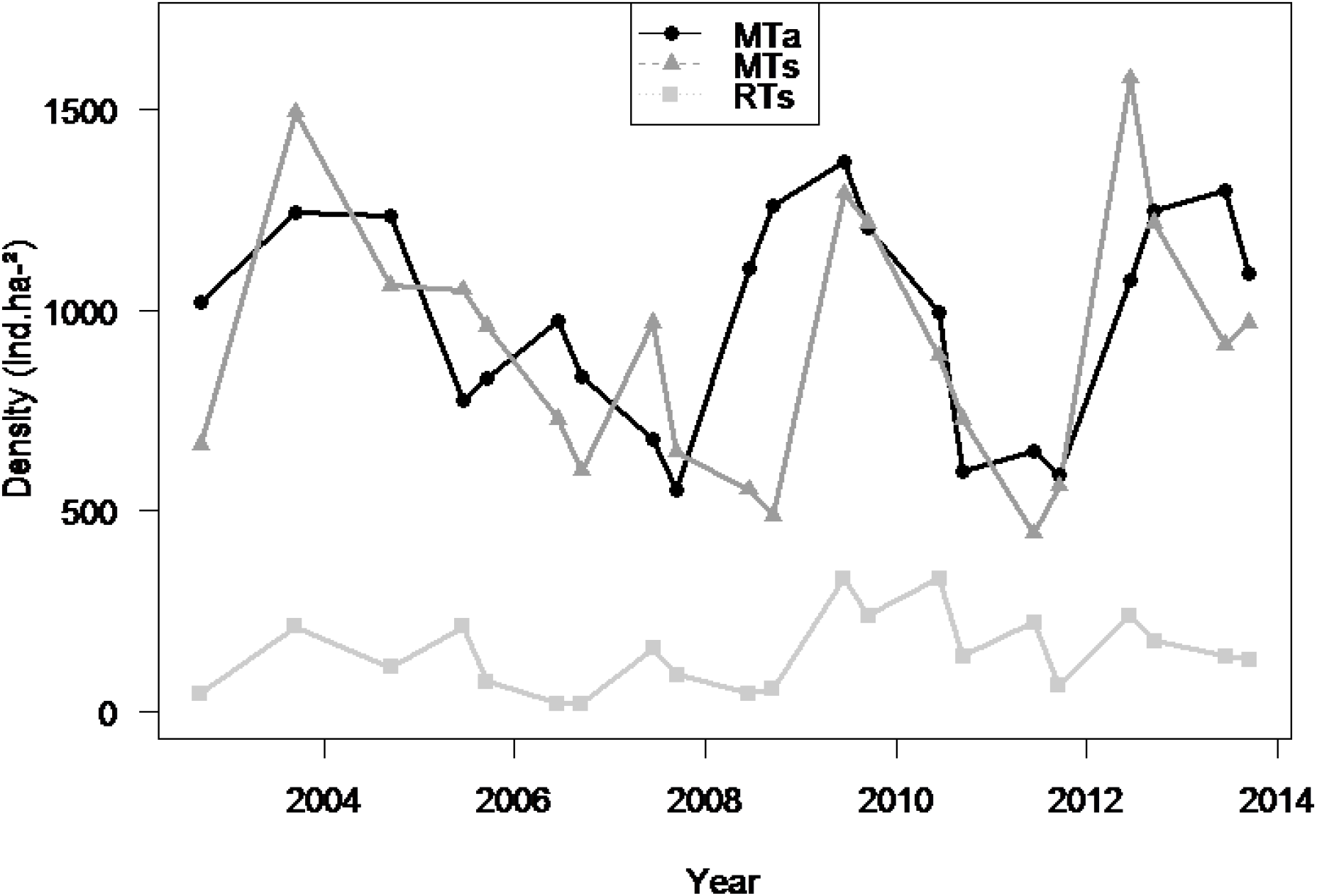
Densities of rainbow trout and marble trout (1+ and older ind · ha^-1^) estimated in September and June from 2002 to 2014. Modified from Vincenzi et al. 2011

**Table 1.** Number of fish and their mean total length in mm (±standard deviation) for each class at each sample d in the stable isotope analysis.

|  | < 200 mm |  | > 200 mm |  | YOY |  |
| --- | --- | --- | --- | --- | --- | --- |
|  | N | TL | N | TL | N | TL |
| ***MTa*** |  |  |  |  |  |  |
| June 12 | 38 | 137 ± 34 | 31 | 250 ± 27 | - | - |
| June 13 | 49 | 142 ± 25 | 28 | 248 ± 24 | - | - |
| Sept 13 | 48 | 153 ± 22 | 31 | 274 ± 28 | 21 | 75 ± 5 |
| ***MTs*** |  |  |  |  |  |  |
| June 12 | 45 | 128 ± 28 | 21 | 285 ± 90 | - | - |
| June 13 | 46 | 147 ± 23 | 13 | 269 ± 55 | - | - |
| Sept 13 | 57 | 149 ± 25 | 21 | 248 ± 29 | 21 | 78 ± 9 |
| ***RTs*** |  |  |  |  |  |  |
| June 12 | 36 | 144 ± 20 | 7 | 280 ± 59 | - | - |
| June 13 | 41 | 141 ± 28 | 6 | 243 ± 45 | - | - |
| Sept 13 | 31 | 160 ± 20 | 22 | 250 ± 43 | 20 | 84 ± 6 |

We sampled freshwater invertebrates using a surber net in three habitats (stones in riffles, litter in pools and blocks) within the two sectors for each sampling sessions. We measured dry weight biomass (mg/m^2^) and compared between sectors, results showed no difference in resource availability between allopatry and sympatry sectors (Online Resource 1). Indexes of evenness, diversity and similarity in benthic invertebrates’ communities showed similar taxonomic compositions in Upper Idrijca and Lower Idrijca (Online Resource 2). We collected aquatic invertebrates for baseline correction in the stable isotope analyses and we found that stable isotope signatures of invertebrates were not significantly different between sectors for each sampling period. Thus, we did not correct fish stable isotope signature in this study (Online Resource 3).

### Stable isotope analysis

Stable isotopes of carbon and nitrogen are powerful tools for investigating the flow of energy material through food webs. δ ^13^C is the indicator of energy source and δ^15^N is the indicator of trophic level (McCutchan et al., 2003; Vanderklift & Ponsard, 2003). Fin samples and invertebrate samples were oven dried for 48h at 60°C and ground into a fine homogenous powder using a mill (Spex Certiprep 6750 Freezer/Mill). Stable isotope ratios of carbon and nitrogen were analyzed in a Carlo Erba NC2500 elemental analyzer coupled to a Thermo Finnigan MAT Delta XP isotope ratio mass spectrometer. All stable isotope analyses were performed at the Cornell Isotope Laboratory, Cornell University, USA.

We estimated differences in niche centroid location (the mean of δ^13^C and δ^15^N of all individuals in the given group) with a residual permutation procedure (RPP, *n* = 9999, Turner et al., 2010) to test for niche shift, where significant results suggest differences in resources use. We coupled niche centroid location analysis with overlap of standard ellipse area within a Bayesian framework (SEA_b_, SIBER, Jackson et al., 2011). The standard ellipse area is a measure of niche breadth, while overlap between SEA_b_ is a proxy of trophic similarity between two groups. First, we assessed the potential competitive interaction between MTs and RTs by estimating SEA_b_ overlaps and niche centroid locations. Then, we investigated the effects of rainbow trout on marble trout resource use by focusing on differences in niche centroid location and SEA_b_ overlaps between MTa and MTs. Finally, we compared position of MTa and RT in the bivariate isotopic plot to evaluate a potential trophic niche replacement of marble trout by rainbow trout in the food web. We quantified niche breadth for each population by calculating SEA_b_ values (‰^2^) and we compared the relative size of ellipses between populations for each sampling period and class to assess niche expansion under sympatry. All of these comparisons in centroid locations and SEA_b_ overlaps were run for each sampling period (June 2012, June 2013 and September 2013) for the three classes and at the population level (i.e. all individuals) for MTa, MTs and RTs.

We assessed differences in trophic position (TP) between MTa and MTs, we converted δ^15^N data for each fish *i* using the formula TP_*i*_ = [(δ^15^N_i_-δ^15^N_baseline_)/3.4] +2 (Anderson & Cabana, 2007). We calculated the average TP of the overall MTa and MTs populations for each sample period and each class and we used t-test to test for significant differences in trophic positions between these two populations. Primary consumer (Heptageniidae) isotopic signatures were used as baseline in the formula.

We used SIBER (Jackson et al., 2011) to test for statistical differences in niche breadth and overlaps estimation All statistical analyses were performed using R (3.1.0 version, R Development Core Team 2012) and ‘siar’ package (Parnell & Jackson, 2013) for stable isotope metrics analysis.

## Results

### Hypothesis 1- Niche segregation in sympatry

In June 2012, centroids niche of MTs and RTs occupied same locations either at the population level or for the two size-classes (Table 2). This result was consistent with substantial SEA_b_ overlaps recorded at the population level (SEA_b_ overlap: 59%), which was to be ascribed mainly to trophic similarity between MTs and RTs < 200-mm (SEA_b_ overlap: 62%, Fig. 2. A, Table 2). In June 2013, centroid locations differed between MTs and RTs for the overall populations and also for the two size classes, which was consistent with the very low overlaps when considering the whole population (SEA_b_ overlap: 5%), only < 200 mm trout (SEA_b_ overlap: 0%, Fig. 2. B) and only > 200 mm (SEA_b_ overlap: 1%, Fig. 3. B). MTs and RTs niche centroid locations differed in September 2013 although we observed a slight overlap (SEA_b_ overlap: 25%) when considering the whole populations (Table 2). Trophic niche breadths [as standard ellipse area (SEA)] were not significantly different between MTs and RTs for whole population, < 200 mm trout and > 200 mm trout, for each sampling periods (Table 2). For YOY, we observed significant differences in niche centroids location, but no overlap and no difference in niche breadths (Fig. 4, Table 2).

**Fig. 2.**
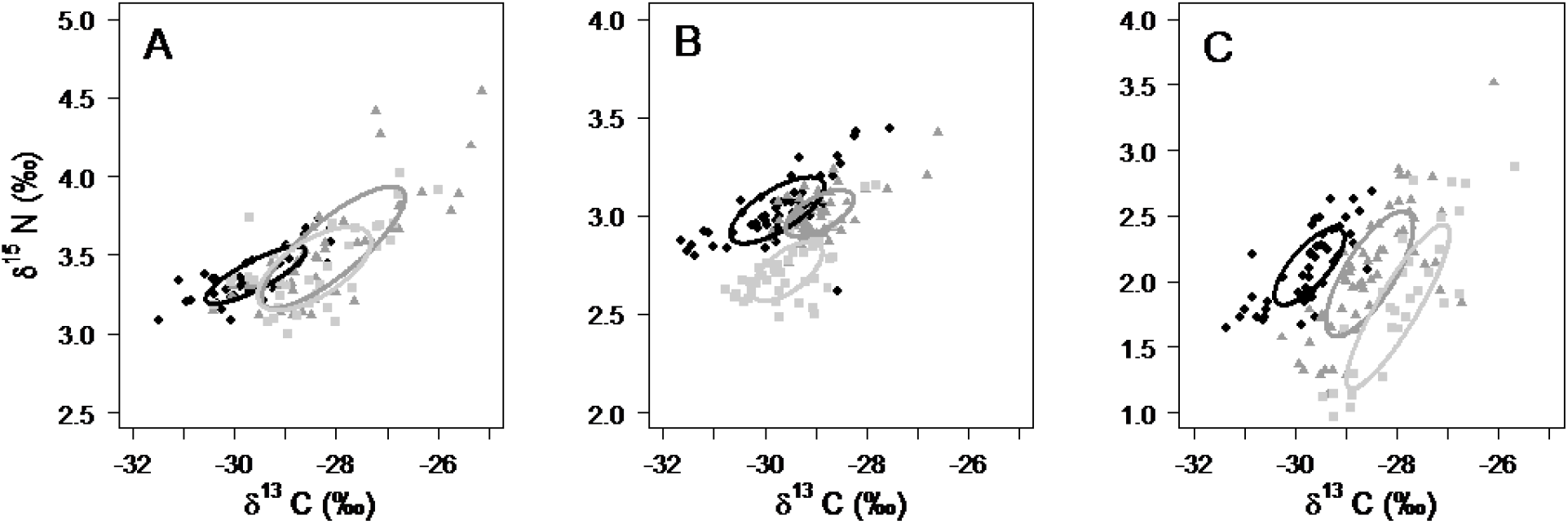
Stable isotope bi-plots of *S. marmoratus* in allopatry (MTa, black circles), in sympatry (MTs, dark grey triangles) and *O. mykiss* in sympatry (RTs, light grey squares) for < 200 mm trout in June 2012 (A), June 2013 (B) and September 2013 (C). Standard ellipses areas (dietary niche breadth) are represented with a solid black line for MTa, dark grey line for MTs and solid light grey for RTs. Note differences in scales on the δ^15^N axis

**Fig. 3.**
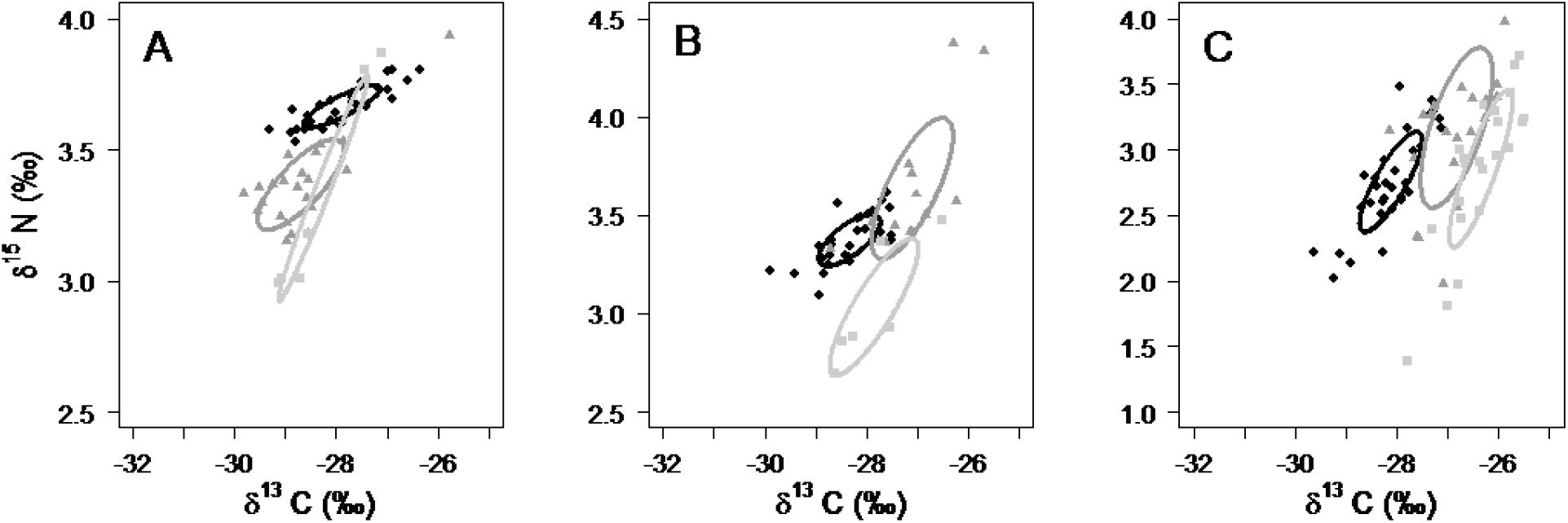
Stable isotope bi-plots of *S. marmoratus* in allopatry (MTa, black circles), in sympatry (MTs, dark grey triangles) and *O. mykiss* in sympatry (RTs, light grey squares) for > 200 mm trout in June 2012 (A), June 2013 (B) and September 2013 (C). Standard ellipses areas (dietary niche breadth) are represented with a solid black line for MTa, dark grey line for MTs and solid light grey for RTs. Note differences in scales on the δ^15^N axis

**Fig. 4.**
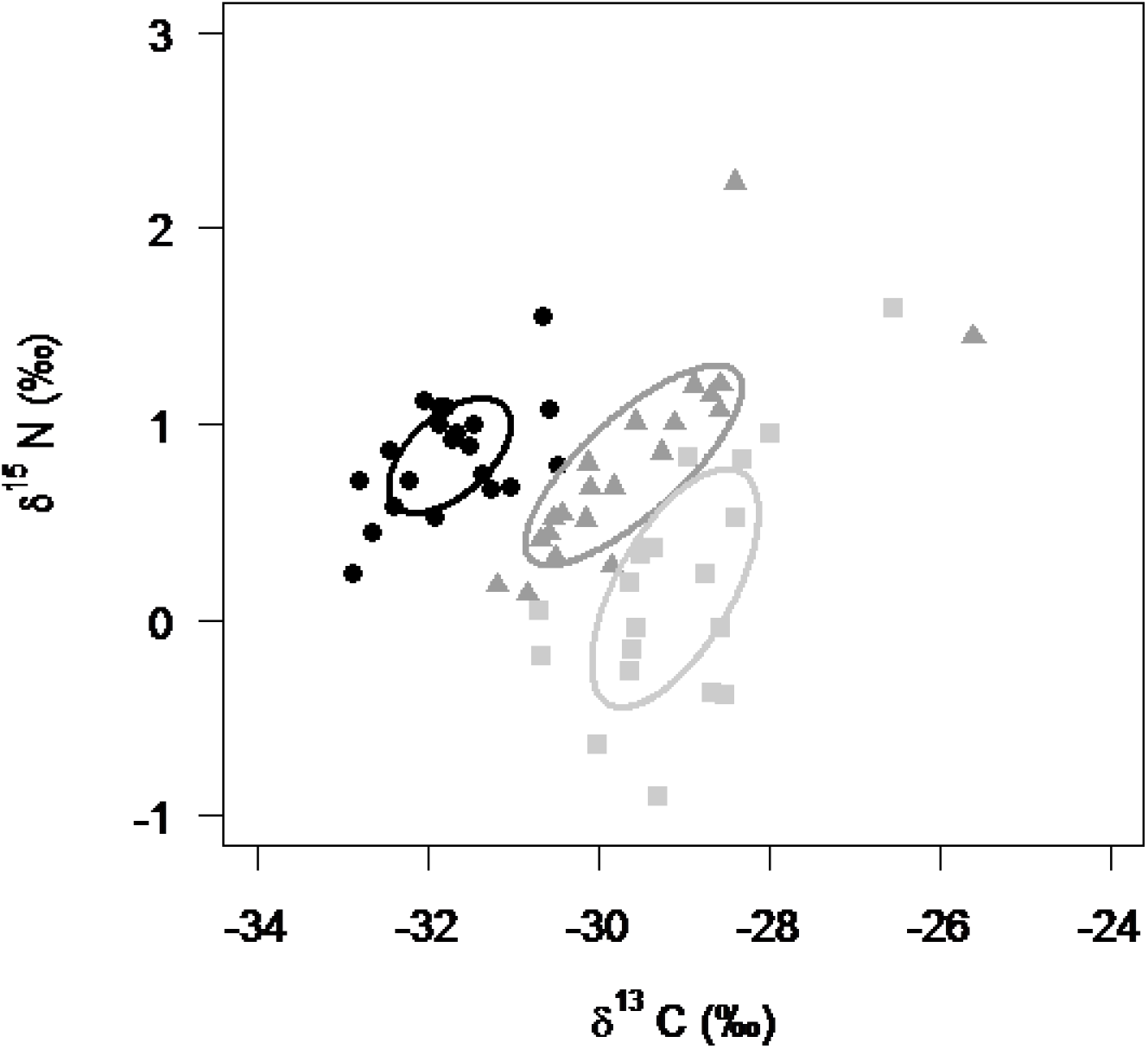
Niche segregation and niche shift on stable isotope bi-plot of YOY *S. marmoratus* in allopatry (MTa, black circles), *S. marmoratus* in sympatry (MTs, dark grey triangles) and *O. mykiss* in sympatry (RTs, light grey squares) with their dietary niche breadth (standard ellipse area) where solid black line is MTa, solid dark grey line is MTs and solid light grey is RTs

**Table 2.** Comparisons of niche centroid locations, niche overlaps(%) and trophic niche breadth (SEA_b_, %^2^) by sampling period for MTs/RTs, MTa/MTs and MTa/RTs 612 comparisons (overall populations and each class). Significant differences in centroid location (α<0.05) are in bold and significant differences in niche breadth are displayed 613 with ^*^, values in parentheses are the ratios MTs SEA_b_/RTs SEA_b_, MTa SEA_b_/MTs SEA _b_ and MTa SEA_b_/RTs SEA_b_ when niche breadths were significantly differen

|  | Sampling period | Centroid location |  |  |  | Niche overlap (%) |  |  |  | Trophic niche breath (SEA<sub>b</sub> %<sup>o</sup>²) |  |  |  |
| --- | --- | --- | --- | --- | --- | --- | --- | --- | --- | --- | --- | --- | --- |
|  |  | Population | <200mm | >200mm | YOY | Population | <200mm | >200mm | YOY | Population | <200mm | >200mm | YOY |
| MTs – RTs |  |  |  |  |  |  |  |  |  |  |  |  |  |
|  | June 12 | 0.52 | 0.18 | 0.23 | - | 59 | 62 | 6 | - | 1.06/1.02 | 1.35/1.10 | 0.95/1.82 | - |
|  | June 13 | <0.01 | <0.001 | <0.01 | - | 5 | 0 | 1 | - | 1.46/1.44 | 0.51/0.57 | 1.29/1.94 | - |
|  | Sept 13 | <0.01 | <0.05 | <0.01 | <0.05 | 25 | 3 | 3 | 0 | 2.08/2.47 | 1.02/1.32 | 1.36/1.07 | 1.78/1.79 |
| MTa – MTs |  |  |  |  |  |  |  |  |  |  |  |  |  |
|  | June 12 | <0.01 | <0.001 | <0.001 | - | 20 | 7 | 0 | - | 0.75/1.06* (1.4) | 0.80 / 1.35* (1.7) | 0.62/0.90 | - |
|  | June 13 | <0.001 | <0.001 | <0.001 | - | 3 | 4 | 1 | - | 0.88/1.46* (1.6) | 0.34/0.19 | 0.60/1.29* (2.1) | - |
|  | Sept 13 | <0.001 | <0.001 | <0.001 | <0.001 | 0 | 0 | 0 | 0 | 1.41/2.07* (1.5) | 0.66/1.02* (1.5) | 0.75/1.36* (1.8) | 0.96/1.78* (1.8) |
| MTa – RTs |  |  |  |  |  |  |  |  |  |  |  |  |  |
|  | June 12 | <0.05 | 0.001 | 0.13 | - | 16 | 8 | 2 | - | 0.75/1.03* (1.4) | 0.80/1.09 | 0.65/1.82* (2.8) | - |
|  | June 13 | <0.001 | <0.05 | <0.05 | - | 0 | 0 | 0 | - | 0.89/1.45* (1.6) | 0.70/0.57 | 0.60/1.93* (3.2) | - |
|  | Sept 13 | <0.001 | <0.001 | <0.001 | <0.01 | 0 | 0 | 0 | 0 | 1.42/2.47* (1.7) | 0.65/1.33* (2.0) | 0.76/1.07 | 0.96/1.78* (1.9) |

### Hypothesis 2- No displacement of the marble trout’s niche

For each class and overall at the population level we observed differences in centroid locations for the three sampling periods (Table 2). Moreover, we observed low overlap between MTa and MTs with the maximum overlap observed in June 2012 at the population level of 20% (Table 2).

Differences in centroid locations and low overlap indicated niche shift between the MTa and MTs for both < 200 mm (Fig. 2), > 200 mm (Fig. 3) and YOY (Fig. 4).

Trophic positions (TPs) of MTa and MTs of overall populations were not significantly different in either in June 2012 or June 2013 (2012: *t* = -0.7, *df* = 108, *p* = 0.94, TP_MTs_=3.87, TP_MTa_=3.87; 2013: *t* = -0.3, *df* = 99, *p* = 0.76, TP_MTs_=3.57, TP_MTa_=3.57). There was no difference in TP in September 2013 ( *t* = 1.3, *df* = 188, *p* = 0.18, TP^MTs^=3.48 and TP^MTa^=3.44). Differences in TP of MTa and MTs for trout < 200 mm were significant in June 2012 ( *t* = 2.2, *df* = 62, *p* < 0.05) with a higher TP for MTs (TP_MTs_=3.89 and TP^MTa^=3.82), but not in June 2013 and September 2013 (June: *t* = -0.1, *df* = 93, *p* = 0.87; *t* = 1.6 _TPMTs_=3.53, TP^MTa^=3.53; Sept: *t* = 2.6, *df* = 95, *p* = 0.10, TP^MTs^=3.49 and TP^MTa^=3.43). For bigger fishes, TP between MTa and MTs were always different with a higher TP for MTa in June 2012 ( *t* = -7.1, *df* = 28, *p* < 0.001, TP^MTs^=3.83 and TP^MTa^=3.91). For both June and September 2013, trophic position of MTs was higher than MTa ( *t* = 2.5, *df* = 13, *p* < 0.05, TP^MTs^=3.71, TP^MTa^=3.63 and *t* = 4.0, *df* = 31, *p* < 0.001, TP^MTs^=3.82, TP^MTa^=3.64, respectively). TPs of MTa YOY and MTs YOY were not significantly different for ( *t* = 1.04, *df* = 32, *p* = 0.30, TP^MTs^=3.12 and TP^MTa^=3.08).

### Hypothesis 3- No niche breadth difference

Across sampling occasions, trophic niche breadth of the MTs whole population was from 1.4 to 1.6 times larger than the MTa trophic niche breadth (Table 2). Niche of MTs trout < 200 mm was significantly larger than MTa nice both in June 2012 and September 2013 (1.7 and 1.5 times, respectively), while niche of MTs > 200 mm was significantly larger both in June 2013 and September 2013 (2.1 and 1.8 times, respectively, Table 2). The niche of MTs YOY was 1.8 greater than MTa YOY during fall 2013 (Fig. 4, Table 2).

### Hypothesis 4- No niche replacement

Niche centroids of MTa and RTs for trout > 200 mm were not different in June 2012 and the overlap was low (Table 2). Niche centroids were significantly different for < 200 mm trout and the whole populations for the same year. For other sampling periods, niche centroids were different and we did not observe overlaps in any class (Table 2). Overall, RTs displayed a larger niche (from 1.4 to 1.7 times) than MTa (Table 2).

## Discussion

We used a trophic niche approach to find the determinants of the stable coexistence of sympatric native and non-native salmonids in a Slovenian stream. We expected separate trophic niches between the two species and no ecological consequence for the marble trout arising from the resource partitioning.

### Hypothesis 1- Niche segregation in sympatry

The hypothesis of niche segregation in sympatry between marble trout and rainbow trout was only partially supported, depending on sampling date. Trophic niches of the two sympatric salmonids overlapped, thus showing sharing of resources, in 2012 for < 200 mm MTs and RTs. This is consistent with previous studies that showed diet similarities between native and non-native stream resident salmonids (Dunham et al., 2000; Hilderbrand & Kershner, 2004; McHugh et al., 2008). In our study, MTs and RTs niches overlapped while the biomass of benthic invertebrates was the highest measured in this study, suggesting sufficient ressource. However, except for the case of < 200 mm trout in June 2012, our first hypothesis was supported by trophic niche segregation between MTs and RTs. Niche overlap between marble and rainbow trout in sympatry ranged from 0 % to 6 %, which is low for taxonomically related fishes living in sympatry. For example, trophic niches of the invasive topmouth gudgeon (*Pseudorasbora parva*) and native roach (*Rutilus rutilus*) and commun rudd (*Scardinus erythropthalmus*) living in ponds in England and Wales overlapped from 49 % to 73 % and from 86 % to 92 %, respectively (Jackson & Britton, 2013). A study of native cyprinids in the Middle East found niche overlap up to 72% between the native *Capoeta damascina* and the endangered *Garra ghorensis* (Hamidan et al., 2015).

Trophic niche segregation between native and non-native salmonids has been often observed in lakes with pelagic and littoral feeding areas (Langeland et al., 1991; Eloranta et al., 2014). In streams, the two different feeding areas for salmonids are pools and riffles; unfortunately, stable isotope signatures of benthic invertebrates living in pools and riffles were too similar to investigate preferences in the use of feeding areas in the two trout species. Resource partitioning for benthic food often enables sympatric native salmonids to coexist. While drift of terrestrial invertebrates represents an ecological opportunity for trout, the reliance of the rainbow trout in its introduced area on terrestrial subsidies is well known (Nakano et al., 1999; Nakano & Murakami, 2001; Baxter et al., 2007). This may explain the niche segregation between MTs and RTs in our study. The isotopic signatures of rainbow trout for September 2013 are less depleted in 13C than marble trout signatures such as isotopic signatures of terrestrial invertebrates when compared to benthic invertebrates.

### Hypothesis 2- No displacement of the marble trout’s niche

Contrary to our predictions, we observed centroids shifts and low trophic niche overlaps (20% maximum) between MTa and MTs for the three size/age classes (YOY, < 200 mm, > 200 mm). These results indicated a shift of the trophic niche of marble trout occurring in sympatry with rainbow trout in each of the three sampling periods. Trophic niche shifts patterns between coexisting native and non-native species often result in niche divergence (Miyasaka et al., 2003; Mason et al., 2005; Tran et al., 2015). Surprisingly, our results showed that diet of MTs became more similar to RTs than MTa, pointing to niche convergence of MTs to RTs. The convergence of niches of native and non-native salmonids has rarely been found. Cucherousset et al. (2007) found niche convergence between native brown trout *Salmo trutta* and non-native brook trout *Salvelinus fontinalis* and proposed behavioural interactions as the mechanism driving the observed convergence. Previous studies have shown the ability of fish to copy (social learning) other individuals to guide their foraging behaviour (Pike & Laland, 2010; Pike et al., 2010).

Trophic positions of marble trout in allopatry ranged from 3.49 to 3.91 and from 3.43 to 3.89 in sympatry, which indicate that marble trout occupies a high trophic level and confirm previous observations of cannibalism in other marble trout populations living in allopatry (Jesenšek and Crivelli, *pers. obs*). Also, these results show that the species occupies similar trophic levels in sympatry with rainbow trout, thus the niche shift is mostly due to an increase of δ ^13^C in trout fins. The increase of δ ^13^C highlights a change in the relative contribution of diet resources (Post, 2002; McCutchan et al., 2003) and this might be due to the addition of rainbow trout as new prey in the diet of marble trout (instead of strict cannibalism in allopatry). Since fish predators usually select conspecifics over fish of other species (Byström et al., 2013), our findings deserve further investigations to distinguish the strict cannibalism of marble trout from the piscivory on rainbow trout.

### Hypothesis 3- No niche breadth difference

We found a broader trophic niche for MTs compared to MTa in all sampling periods, which suggest trophic niche expansion for marble trout when living in sympatry with rainbow trout. In September 2013, YOY of MTs displayed a niche almost twice as big as that of MTa YOY. Niche size of MTa > 200 mm were 1.5 times larger than that of MTs < 200 mm. Contrary to what is typically observed, i.e. size of trophic niches that is limited by competition with sympatric species (Pianka, 1974; Codron et al., 2011; Hamidan et al., 2015), our results showed that size of niches of marble trout when sympatry with rainbow trout were wider than when in allopatry. Interspecific competition in closely related species might have effects similar to those of intraspecific competition. Increase in intraspecific competition (e.g. higher density) leads to a reduction of the availability of the preferred prey and to an increase in diversity of resources (Svanbäck & Bolnick, 2007; Araújo et al., 2011), and similarly interspecific competition between marble trout and rainbow trout increased the trophic niche of the native species. Moreover, Musseau et al. (2015) have shown that in marble trout an increase in density of <200 mm fish led to an increase of trophic niche breadth, which is consistent with observed overlap of 62% between < 200 mm trout MTs and RTs. However, since – as we explain below - rainbow trout did not use trophic subsidies of the marble trout, marble trout was not forced by the non-native species to prey on unsuitable invertebrates.

### Hypothesis 4- No niche replacement

We found a trophic shift by native marble trout from allopatry to sympatry but not a niche replacement since the trophic niche of non-native rainbow trout did not occupy the trophic niche of the native marble trout in allopatry (niche overlap from 0% to 16%). Rainbow trout is known to take food from other salmonids (Baxter et al., 2007), but in our case the trophic shift we observed in marble trout in sympatry with rainbow trout was not caused by trophic replacement by rainbow trout. The absence of trophic replacement suggests that the two salmonids have different functional traits as previously shown in other studies (Baxter et al., 2007; Benjamin et al., 2013).

### Conclusions and perspectives

Our results showed unexpected ecological consequences of non-native rainbow trout on native marble trout. Rainbow trout caused a shift in the trophic niche of native marble trout; however, this diet shift only minor effects on survival, growth, or fish density of marble trout (Vincenzi et al., 2011). Density of rainbow trout in sympatry with marble trout was stable over time and much lower (143 ± 91 ind·ha^-1^ between 2002-2014) than density of rainbow trout when occurring in sympatry with hybrid trout (3 populations, Kozjak: 550 ± 243.8 ind·ha^-1^ between 2008-2013; Dabreck: 750 ± 388.2 ind·ha^-1^ between 2008-2014 and Hotenja: 428 ± 267.8 ind·ha^-1^ between 2008-2014) and of the rainbow trout population living in allopatry in the Soča basin (Godiča: 1290.6 ± 813.4 ind·ha^-1^ between 2010 and 2014). The large differences in density suggest that the trophic interactions between hybrid trout and rainbow trout might be different than those between marble trout and rainbow trout and that native marble trout exerts a control on rainbow trout.

Further study on diet composition and piscivory rate along with prey identification are necessary to disentangle piscivory from strict cannibalism in marble trout. Stable isotope analysis cannot discriminate between species, but barcoding analysis of stomach content or faeces may help identify the direction of predation (marble on rainbow or *viceversa*) (Parsons et al., 2005).

## Acknowledgements

This study has been mostly funded by the MAVA Foundation and the Tolmin Angling Association. We warmly thank the field team of the Tolmin Angling Association for the hard work carried out in a friendly atmosphere and Isabel Cantera for her helpful participation during the fieldwork. We are grateful to the two anonymous reviewers for their valuable comments on earlier version of this manuscript. The authors declare that they have no conflict of interest.

## References

1. Anderson, C., & G. Cabana, 2007. Estimating the trophic position of aquatic consumers in river food webs using stable nitrogen isotopes. Journal of North American Benthological Society 26: 273–285.

2. Araújo, M. S., D. I. Bolnick, & C. A. Layman, 2011. The ecological causes of individual specialisation. Ecology Letters 14: 948–958, http://onlinelibrary.wiley.com/doi/10.1111/j.1461-0248.2011.01662.x/full.

3. Baxter, C. V., K. D. Fausch, M. Murakami, & P. L. Chapman, 2007. Invading rainbow trout usurp a terrestrial prey subsidy from native charr and reduce their growth and abundance. Oecologia 153: 461–470.

4. Benjamin, J. R., & C. V. Baxter, 2012. Is a trout a trout? A range-wide comparison shows nonnative brook trout exhibit greater density, biomass, and production than native inland cutthroat trout. Biological Invasions 14: 1865–1879.

5. Benjamin, J. R., F. Lepori, C. V. Baxter, & K. D. Fausch, 2013. Can replacement of native by non-native trout alter stream-riparian food webs?. Freshwater Biology 58: 1694–1709.

6. Berrebi, P., M. Povz, D. Jesensek, G. Cattaneo-berrebi, & A. J. Crivelli, 2000. The genetic diversity of native, stocked and hybrid populations of marble trout in the Soca river, Slovenia. Heredity 85: 277–287.

7. Blanchet, S., G. Loot, G. Grenouillet, & S. Brosse, 2007. Competitive interactions between native and exotic salmonids: a combined field and laboratory demonstration. Ecology of Freshwater Fish 16: 133–143, http://doi.wiley.com/10.1111/j.1600-0633.2006.00205.x.

8. Bøhn, T., P.-A. Amundsen, & A. Sparrow, 2008. Competitive exclusion after invasion?. Biological Invasions 10: 359–368.

9. Byström, P., P. Ask, J. Andersson, & L. Persson, 2013. Preference for cannibalism and ontogenetic constraints in competitive ability of piscivorous top predators. PLoS ONE 8: e70404.

10. Carlson, S. M., A. P. Hendry, & B. H. Letcher, 2007. Growth rate differences between resident native brook trout and non-native brown trout. Journal of Fish Biology 71: 1430–1447.

11. Chesson, P., 2000. Mechanisms of maintenance of species diversity. Annual Review of Ecology and Systematics 31: 343–366.

12. Clavero, M., & E. García-Berthou, 2005. Invasive species are a leading cause of animal extinctions. Trends in Ecology and Evolution 20: 110, http://www.ncbi.nlm.nih.gov/pubmed/16701353.

13. Codron, D., J. Hull, J. S. Brink, J. Codron, D. Ward, & M. Clauss, 2011. Effect of competition on niche dynamics of syntopic grazing ungulates: contrasting the predictions of habitat selection models using stable isotope analysis. Evolutionary Ecology Research 13: 217–235.

14. Cucherousset, J., J.-C. Aymes, F. Santoul, & R. Céréghino, 2007. Stable isotope evidence of trophic interactions between introduced brook trout Salvelinus fontinalis and native brown trout Salmo trutta in a mountain stream of south-west France. Journal of Fish Biology 71: 210–223, http://doi.wiley.com/10.1111/j.1095-8649.2007.01675.x.

15. Cucherousset, J., & J. D. Olden, 2011. Ecological impacts on non-native freshwater fishes. Fisheries 36: 215–230.

16. Dunham, J. B., M. E. Rahn, R. E. Schroeter, & S. W. Breck, 2000. Diets of sympatric Lahontan cutthroat trout and nonnative brook trout: Implications for species interactions. Western North American Naturalist 60: 304–310.

17. Eloranta, A. P., P. Nieminen, & K. K. Kahilainen, 2014. Trophic interactions between introduced lake trout (Salvelinus namaycush) and native Arctic charr (S. alpinus) in a large Fennoscandian subarctic lake. Ecology of Freshwater Fish 24: 181–192.

18. Fumagalli, L., A. Snoj, D. Jesenšek, F. Balloux, T. Jug, O. Duron, F. Brossier, A. J. Crivelli, & P. Berrebi, 2002. Extreme genetic differentiation among the remnant populations of marble trout (Salmo marmoratus) in Slovenia. Molecular Ecology 11: 2711–2716.

19. Hamidan, N., M. C. Jackson, & J. R. Britton, 2015. Diet and trophic niche of the endangered fish Garra ghorensis in three Jordanian populations. Ecology of Freshwater Fish 1–10, http://doi.wiley.com/10.1111/eff.12226.

20. Hasegawa, K., N. Ishiyama, & H. Kawai, 2014. Replacement of nonnative rainbow trout by nonnative brown trout in the Chitose River system, Hokkaido, northern Japan. Aquatic Invasions 9: 221–226, http://www.aquaticinvasions.net/2014/issue2.html.

21. Hasegawa, K., T. Yamamoto, & S. Kitanishi, 2010. Habitat niche separation of the nonnative rainbow trout and native masu salmon in the Atsuta River, Hokkaido, Japan. Fisheries Science 76: 251–256.

22. Hasegawa, K., T. Yamamoto, M. Murakami, & K. Maekawa, 2004. Comparison of competitive ability between native and introduced salmonids: Evidence from pairwise contests. Ichthyological Research 51: 191–194.

23. Hasegawa, K., C. Yamazaki, T. Ohta, & K. Ohkuma, 2012. Food habits of introduced brown trout and native masu salmon are influenced by seasonal and locational prey availability. Fisheries Science 78: 1163–1171.

24. Hilderbrand, R. H., & J. L. Kershner, 2004. Influence of habitat type on food supply, selectivity, and diet overlap of Bonneville cutthroat trout and nonnative brook trout in Beaver Creek, Idaho. North American Journal of Fisheries Management 24: 33–40, http://www.tandfonline.com/doi/abs/10.1577/M02-192.

25. Holt, R. D., 2009. Bringing the Hutchinsonian niche into the 21st century: ecological and evolutionary perspectives. Proceedings of the National Academy of Sciences of the United States of America 106: 19659–19665, http://www.pubmedcentral.nih.gov/articlerender.fcgi?artid=2780934&tool=pmcentrez&rendertype=abstract.

26. Houde, A. L. S., C. C. Wilson, & B. D. Neff, 2015a. Effects of competition with four nonnative salmonid species on Atlantic salmon from three populations. Transactions of the American Fisheries Society 144: 1081–1090.

27. Houde, A. L. S., C. C. Wilson, & B. D. Neff, 2015b. Competitive interactions among multiple non-native salmonids and two populations of Atlantic salmon. Ecology of Freshwater Fish 24: 44–55, <Go to ISI>://000346347100005.

28. Hutchinson, G. E., 1957. Concluding remarks. Cold Spring Symposium on Quantitative biology 22: 415–427, http://www.ncbi.nlm.nih.gov/pubmed/24297239.

29. Inoue, M., H. Miyata, Y. Tange, & Y. Taniguchi, 2009. Rainbow trout (Oncorhynchus mykiss) invasion in Hokkaido streams, northern Japan, in relation to flow variability and biotic interactions. Canadian Journal of Fisheries and Aquatic Sciences 66: 1423–1434.

30. Jackson, A. L., R. Inger, A. C. Parnell, & S. Bearhop, 2011. Comparing isotopic niche widths among and within communities: SIBER - Stable Isotope Bayesian Ellipses in R. Journal of Animal Ecology 80: 595–602.

31. Jackson, M. C., & J. R. Britton, 2013. Variation in the trophic overlap of invasive Pseudorasbora parva and sympatric cyprinid fishes. Ecology of Freshwater Fish 22: 654–657, http://doi.wiley.com/10.1111/eff.12063.

32. Jardine, T. D., M. A. Gray, S. M. McWilliam, & R. A. Cunjak, 2005. Stable isotope variability in tissues of temperate stream fishes. Transactions of the American Fisheries Society 134: 1103–1110.

33. Juncos, R., D. Milano, P. J. Macchi, & P. H. Vigliano, 2015. Niche segregation facilitates coexistence between native and introduced fishes in a deep Patagonian lake. Hydrobiologia 747: 53–67, http://link.springer.com/10.1007/s10750-014-2122-z.

34. Korsu, K., A. Huusko, & T. Muotka, 2009. Invasion of north European streams by brook trout: hostile takeover or pre-adapted habitat niche segregation?. Biological Invasions 12: 1363–1375, http://link.springer.com/10.1007/s10530-009-9553-x.

35. Krueger, C. C., & B. May, 1991. Ecological and genetic effects of salmonid introductions. Canadian Journal of Fisheries and Aquatic Sciences 48: 66–77.

36. Langeland, A., J. H. L’Abee-Lund, B. Jonsson, & N. Jonsson, 1991. Resource partitioning and niche shift in Artic charr Salvelinus alpinus and brown trout Salmo trutta. Journal of Animal Ecology 60: 895–912.

37. Layman, C. A., M. S. Araújo, R. Boucek, C. M. Hammerschlag-Peyer, E. Harrison, Z. R. Jud, P. Matich, A. E. Rosenblatt, J. J. Vaudo, L. a Yeager, david M. Post, & S. Bearhop, 2012. Applying stable isotopes to examine food-web structure: an overview of analytical tools. Biological Reviews 87: 542–562, http://onlinelibrary.wiley.com/doi/10.1111/j.1469-185X.2011.00208.x/full.

38. Layman, C. A., D. A. Arrington, C. G. Montaña, & D. M. Post, 2007. Can stable isotope ratios provide for community-wide measures of trophic structure? Comment. Ecology 88: 42–48, http://www.ncbi.nlm.nih.gov/pubmed/18724745.

39. Lowe, S., B. M., S. Boudjelas, & M. De Poorter, 2000. 100 of the World’s worst invasive alien species. Published by The Invasive Species Specialist Group (ISSG), SSC, IUCN, http://interface.creative.auckland.ac.nz/database/species/reference_files/100English.pdf.

40. Macarthur, R., & R. Levins, 1967. The limiting similarity, convergence, and divergence of coexisting species. The American Naturalist 101: 377–385.

41. Mack, R. N., C. D. Simberloff, W. M. Lonsdale, H. Evans, M. Clout, F. A. Bazzaz, D. Simberloff, W. M. Lonsdale, H. Evans, M. Clout, & F. A. Bazzaz, 2000. Biotic invasions: causes, epidemiology, global consequences, and control. Ecological Applications 10: 689–710.

42. Mason, N., D. Mouillot, W. Lee, & J. Wilson, 2005. Functional richness, functional evenness and functional divergence: the primary components of functional diversity. Oikos 111: 112–118, http://onlinelibrary.wiley.com/doi/10.1111/j.0030-1299.2005.13886.x/full.

43. Mason, N. W. H., P. Irz, C. Lanoiselée, D. Mouillot, & C. Argillier, 2008. Evidence that niche specialization explains species-energy relationships in lake fish communities. The Journal of animal ecology 77: 285–296, http://www.ncbi.nlm.nih.gov/pubmed/18179548.

44. McCutchan, J. H., W. M. Lewis, C. Kendall, & C. C. Mcgrath, 2003. Variation in trophic shift for stable isotope ratios of carbon, nitrogen, and sulfur. Oikos 102: 378–390.

45. McHugh, P., P. Budy, G. Thiede, & E. Vandyke, 2008. Trophic relationships of nonnative brown trout, Salmo *trutta*, and native Bonneville cutthroat trout, *Oncorhynchus clarkii utah*, in a northern Utah, USA river. Environmental Biology of Fishes 81: 63–75.

46. Miyasaka, H., S. Nakano, & T. Furukawa-Tanaka, 2003. Food habit divergence between white-spotted charr and masu salmon in Japanese mountain streams: circumstantial evidence for competition. Limnology 4: 1–10.

47. Morita, K., J.-I.I. Tsuboi, & H. Matsuda, 2004. The impact of exotic trout on native charr in a Japanese stream. Journal of Applied Ecology 41: 962–972, http://doi.wiley.com/10.1111/j.0021-8901.2004.00927.x.

48. Musseau, C., S. Vincenzi, D. Jesensek, I. Cantera, S. Boulêtreau, F. Santoul, & A. J. Crivelli, 2015. Direct and indirect effects of environmental factors on dietary niches in size-structured populations of a wild salmonid. Ecosphere 6: 1–15.

49. Nakano, S., Y. Kawaguchi, Y. Taniguchi, H. Miyasaka, Y. Shibata, H. Urabe, & N. Kuhara, 1999. Selective foraging on terrestrial invertebrates by rainbow trout in a forested headwater stream in northern Japan. Ecological Research 14: 351–360.

50. Nakano, S., & M. Murakami, 2001. Reciprocal subsidies: dynamic interdependence. Proceedings of the National Academy of Sciences 98: 166–170.

51. Newsome, S., C. Martinez del Rio, S. Bearhop, & D. L. Phillips, 2007. A niche for isotopic ecology. Frontiers in Ecology 5: 429–436, http://www.esajournals.org/doi/abs/10.1890/060150.1.

52. Newsome, S., J. Yeakel, P. V. Wheatley, & M. T. Tinker, 2012. Tools for quantifying isotopic niche space and dietary variation at the individual and population level. Journal of Mammalogy 93: 329–341, http://asmjournals.org/doi/abs/10.1644/11-MAMM-S-187.1.

53. Nomoto, K., H. Omiya, T. Sugimoto, K. Akiba, K. Edo, & S. Higashi, 2010. Potential negative impacts of introduced rainbow trout on endangered Sakhalin taimen through redd disturbance in an agricultural stream, eastern Hokkaido. Ecology of Freshwater Fish 19: 116–126.

54. Okabe, F., & N. Agetsuma, 2007. Habitat use by introduced raccoons and native raccoon dogs in a deciduous forest of Japan. Journal of Mammalogy 88: 1090–1097.

55. Parnell, A. C., & A. Jackson, 2013. SIAR: Stable Isotope Analysis in R. R package version 4.2., http://w.download.idg.pl/CRAN/web/packages/siar/siar.pdf.

56. Parsons, K. M., S. B. Piertney, S. J. Middlemas, P. S. Hammond, & J. D. Armstrong, 2005. DNA-based identification of salmonid prey species in seal faeces. Journal of Zoology 266: 275–281, http://doi.wiley.com/10.1017/S0952836905006904.

57. Pearman, P. B., A. Guisan, O. Broennimann, & C. F. Randin, 2008. Niche dynamics in space and time. Trends in Ecology and Evolution 23: 149–158.

58. Pianka, E. R., 1974. Niche overlap and diffuse competition. Proceedings of the National Academy of Sciences of the United States of America 71: 2141–2145.

59. Pike, T. W., J. R. Kendal, L. E. Rendell, & K. N. Laland, 2010. Learning by proportional observation in a species of fish. Behavioral Ecology 21: 570–575.

60. Pike, T. W., & K. N. Laland, 2010. Conformist learning in nine-spined sticklebacks’ foraging decisions. Biology letters 6: 466–468.

61. Post, D. M., 2002. The long and short of food-chain length. Trends in Ecology and Evolution 17: 269–277, http://linkinghub.elsevier.com/retrieve/pii/S0169534702024552.

62. Ricciardi, A., M. F. Hoopes, M. P. Marchetti, & J. L. Lockwood, 2013. Progress toward understanding the ecological impacts of nonnative species. Ecological Monographs 83: 263–282.

63. Sagouis, A., J. Cucherousset, S. Villéger, F. Santoul, & S. Boulêtreau, 2015. Non-native species modify the isotopic structure of freshwater fish communities across the globe. Ecography in press, http://doi.wiley.com/10.1111/ecog.01348.

64. Schoener, T. W., 1974. Resource partitioning in ecological communities. Science 185: 27–39.

65. Seiler, S. M., & E. R. Keeley, 2009. Competition between native and introduced salmonid fishes: cutthroat trout have lower growth rate in the presence of cutthroat–rainbow trout hybrids. Canadian Journal of Fisheries and Aquatic Sciences 66: 133–141, http://www.nrcresearchpress.com/doi/abs/10.1139/F08-194.

66. Stanković, D., A. J. Crivelli, & A. Snoj, 2015. Rainbow trout in Europe: introduction, naturalization, and impacts. Reviews in Fisheries Science & Aquaculture 23: 39–71, http://www.tandfonline.com/doi/full/10.1080/23308249.2015.1024825.

67. Strayer, D. L., 2010. Alien species in fresh waters: ecological effects, interactions with other stressors, and prospects for the future. Freshwater Biology 55: 152–174, http://doi.wiley.com/10.1111/j.1365-2427.2009.02380.x.

68. Svanbäck, R., & D. I. Bolnick, 2007. Intraspecific competition drives increased resource use diversity within a natural population. Proceedings of the Royal Society B: Biological Sciences 274: 839–844, http://www.pubmedcentral.nih.gov/articlerender.fcgi?artid=2093969&tool=pmcentrez&rendertype=abstract.

69. Takami, T., T. Yoshihara, Y. Miyakoshi, & R. Kuwabara, 2002. Replacement of white-spotted charr Salvelinus leucomaenis by brown trout Salmo trutta in a branch of the Chitose River, Hokkaido. Nippon Suisan Gakkaishi 68: 24–28.

70. Tran, T. N. Q., M. C. Jackson, D. Sheath, H. Verreycken, & J. R. Britton, 2015. Patterns of trophic niche divergence between invasive and native fishes in wild communities are predictable from mesocosm studies. Journal of Animal Ecology in press, http://doi.wiley.com/10.1111/1365-2656.12360.

71. Turner, T. F., M. Collyer, & T. J. Krabbenhoft, 2010. A general hypothesis-testing framework for stable isotope ratios in ecological studies. Ecology 91: 2227–2233, http://www.esajournals.org/doi/abs/10.1890/09-1454.1.

72. Van Deventer, J. S., & W. S. Platts, 1989. Microcomputer software system for generating population statistics from electrofishing data-users guide for Microfish 3.0. U.S. Forest Service General Technical Report INT-254.

73. Van Zwol, J. A., B. D. Neff, & C. C. Wilson, 2012. The Effect of Nonnative Salmonids on Social Dominance and Growth of Juvenile Atlantic Salmon. Transactions of the American Fisheries Society 141: 907–918, <Go to ISI>://WOS:000306462500005.

74. Vanderklift, M. A., & S. Ponsard, 2003. Sources of variation in consumer-diet15N enrichment: a meta-analysis. Oecologia 136: 169–182, http://www.ncbi.nlm.nih.gov/pubmed/12802678.

75. Vincenzi, S., A. J. Crivelli, D. Jesensek, & G. A. De Leo, 2008. The role of density-dependent individual growth in the persistence of freshwater salmonid populations. Oecologia 156: 523–534.

76. Vincenzi, S., A. J. Crivelli, D. Jesensek, G. Rossi, & G. A. De Leo, 2011. Innocent until proven guilty? Stable coexistence of alien rainbow trout and native marble trout in a Slovenian stream. Naturwissenschaften 98: 57–66.

77. Wauters, L. A., & J. Gurnell, 1999. The mechanism of replacement of red squirrels by grey squirrels: a test of the interference competition hypothesis. Ethology 105: 1053–1071.

